# Disrupted dynamic functional network connectivity among cognitive control networks in the progression of Alzheimer’s disease

**DOI:** 10.1101/2020.12.31.424877

**Authors:** Mohammad S. E. Sendi, Elaheh Zendehrouh, Zening Fu, Jingyu Liu, Yuhui Du, Elizabeth Mormino, David H. Salat, Vince D. Calhoun, Robyn. L. Miller

## Abstract

**Background:** Alzheimer’s disease (AD) is the most common age-related dementia that promotes a decline in memory, thinking, and social skills. The initial stages of dementia can be associated with mild symptoms, and symptom progression to a more severe state is heterogeneous across patients. Recent work has demonstrated the potential for functional network mapping to assist in the prediction of symptomatic progression. However, this work has primarily used static functional connectivity (sFC) from rs-fMRI. Recently, dynamic functional connectivity (dFC) has been recognized as a powerful advance in functional connectivity methodology to differentiate brain network dynamics between healthy and diseased populations.

**Methods:** Group independent component analysis was applied to extract 17 components within the cognitive control network (CCN) from 1385 individuals across varying stages of AD symptomology. We estimated dFC among 17 components within the CCN, followed by clustering the dFCs into 3 recurring brain states and then estimated a hidden Markov model and the occupancy rate for each subject. Finally, we investigated the link between CCN dFC connectivity features with AD progression.

**Results:** Progression of AD symptoms were associated with increases in connectivity within the middle frontal gyrus. Also, the AD with mild and severer symptoms showed less connectivity within the inferior parietal lobule and between this region with the rest of CCN. Finally, comparing with mild dementia, we found that the normal brain spends significantly more time in a state with lower within middle frontal gyrus connectivity and higher connectivity between the hippocampus and the rest of CCN, highlighting the importance of assessing the dynamics of brain connectivity in this disease.

**Conclusion:** Our results suggest that AD progress not only alters the CCN connectivity strength but also changes the temporal properties in this brain network. This suggests the temporal and spatial pattern of CCN as a biomarker that differentiates different stages of AD.

**Impact Statement:** By assuming that functional connectivity is static over time, many of previous studies have ignored the brain dynamic in Alzheimer’s disease progression. Here, a longitudinal resting-state functional magnetic resonance imaging data are used to explore the temporal changes of functional connectivity in the cognitive control network in Alzheimer’s disease progression. The result of this study would increase our understanding about the underlying mechanisms of Alzheimer’s Disease and help in finding future treatment of this neurological disorder.

## 1 Introduction

Alzheimer’s disease (AD) is the most common age-related dementia, which affects 10-30% of individuals over 65 years of age (Masters et al., 2015). It usually promotes a decline in memory, cognition, everyday function, and social skills. AD usually progresses slowly in 3 stages, including mild cognitive impairment (early-stage), mild dementia (middle-stage), and severe dementia (late-stage) (Ryan and Rossor, 2011). To date, there is no treatment for AD, but some interventions, including pharmacological (Massoud and Léger, 2011) and non-pharmacological (Shigihara et al., 2020; Zucchella et al., 2018) can decelerate its progress in particular when it is detected at an early stage(Yiannopoulou and Papageorgiou, 2020). Predicting AD progression and differentiating different stages of this disease are thus essential steps in early medical intervention for this mental disorder (Badhwar et al., 2017; Brand et al., 2019; Gupta et al., 2019; Kruthika et al., 2019; Lee, Garam, 2019; Z. Wang et al., 2019). Also, knowing that the initial stages of dementia show heterogeneous symptoms across patients, identifying individuals at risk for progression from mild cognitive impairment (MCI) to early or late dementia is challenging (Komarova and Thalhauser, 2011).

In recent years, functional connectivity (FC) obtained from resting-state functional magnetic resonance imaging (rs-fMRI) has shown sensitivity to the prediction of current and future AD based on brain network connectivity (Badhwar et al., 2017; Brier et al., 2014; Wang et al., 2007; Z. Wang et al., 2019; Zhao et al., 2020). A large body of the AD-related literature has focused on default mode network (DMN) FC disruption, and other brain networks are underexplored (Agosta et al., 2012; Balthazar et al., 2014; Miao et al., 2011; Zhong et al., 2014). But other brain networks such as the cognitive control network (CCN) are rarely investigated in AD progression. As previous studies showed, CCNwhich includes inferior parietal lobule, inferior frontal gyrus, middle frontal gyrus, hippocampus, insula, middle cingula cortex, and superior frontal gyrus, is involved in executive function including working memory, internal and external navigation, and attention (Breukelaar et al., 2017; Dhanjal and Wise, 2014; Han et al., 2018; Y. Wang et al., 2019; Westerhausen et al., 2010), which are impaired in AD (Masters et al., 2015). For example, a recent study showed that AD patients have more activation in superior frontal gyrus, middle frontal gyrus, and parietal region compared with normal subjects during a working memory task (Yetkin et al., 2006). Another one found changes in the inferior parietal lobule thickness by progression from normal to MCI (Greene and Killiany, 2010). Another study showed a disconnection between the hippocampus and other brain regions in AD patients (Allen et al., 2007). Therefore, we hypothesized that studying this brain network would reveal useful information about the fundamental neuronal mechanism of AD and its longitudinal progression.

Unlike conventional static FC (sFC), which represents the averaged brain connections over an entire scan, dynamic FC (dFC) refers to brain connectivity within sub-intervals of the time series (Calhoun et al., 2014). In recent years, dFC from rs_fMRI time series has proven highly informative regarding the underlying brain connectivity patterns in different neurological disorders, including schizophrenia (Dong et al., 2019; Sendi et al., 2020a, 2020b), major depressive disorder (Zendehrouh et al., 2020), and AD (Fiorenzato et al., 2019; Fu et al., 2019). Furthermore, features obtained from dFC are shown to be more sensitive to brain disorders, for example, in classifying healthy subjects from patients, than their sFC counterparts (Rashid et al., 2016; Vergara et al., 2018). Any cognitive deficits and clinical symptoms associated with brain disorders likely depend not only on the strength of the connectivity between any pair of the specific brain regions but also on the patterns of temporal variation in the connectivity of those regions.

In this work, from longitudinal rs-fMRI data (LaMontagne et al., 2019), we predicted that CCN dFC in AD and its correlation with behavioral scores could elucidate the pathophysiological mechanism of this neurological disorder. More specifically, we hypothesized that cognitive states in different stages of AD could be linked to the temporal fluctuations exhibited by CCN dFC in this group of subjects. To find this link, we leveraged the sliding window approach followed by k-means clustering to identify a set of connectivity states to investigate dFC within the CCN (Allen et al., 2014). We explored the link between symptom severity in AD with state-specific CCN FC. To further investigate and model the temporal changes in dFC, we estimated the occupancy rate (OCR) for each subject from dFC. Next, we explored the link between these features and cognitive scores via statistical analysis on the estimated OCR features.

## 2 Methods

In this study, ethical approval was granted by the relevant ethics committees, and informed consent was obtained from each subject prior to scanning according to the institutional review board of Washington University School of Medicine.

### Participant

The data is from the Open Access Series of Imaging Studies (OASIS)-3 cohort, which contains imaging and related clinical data of 1098 participants. The data are collected across several ongoing studies in the Washington University Knight Alzheimer Disease Research Center over 15 years. This study used 1385 rs-fMRI and clinical and demographic information at scanning (from 910 subjects) with the age range of 42 to 95 years (LaMontagne et al., 2019). The clinical dementia rating scale sum of boxes (CDR-SOB) scores is used to assess the cognitive stage of the participant at the time of scanning. All subjects must have CDR≤1 at the time of the clinical core assessment and once the participant reached CDR=2 or CDR-SOB>9, they were no longer eligible for the study (LaMontagne et al., 2019). Using CDR-SOB, we categorized all subjects into two groups, including healthy control or HC(CDR-SOB=0), and very mild AD or vmAD (CDR-SOB>0) (O’Bryant et al., 2008). In total, we had 1028 scans of HC, 357 scans of vmAD patients. The demographic information is provided in Table 1.

**Table 1.**
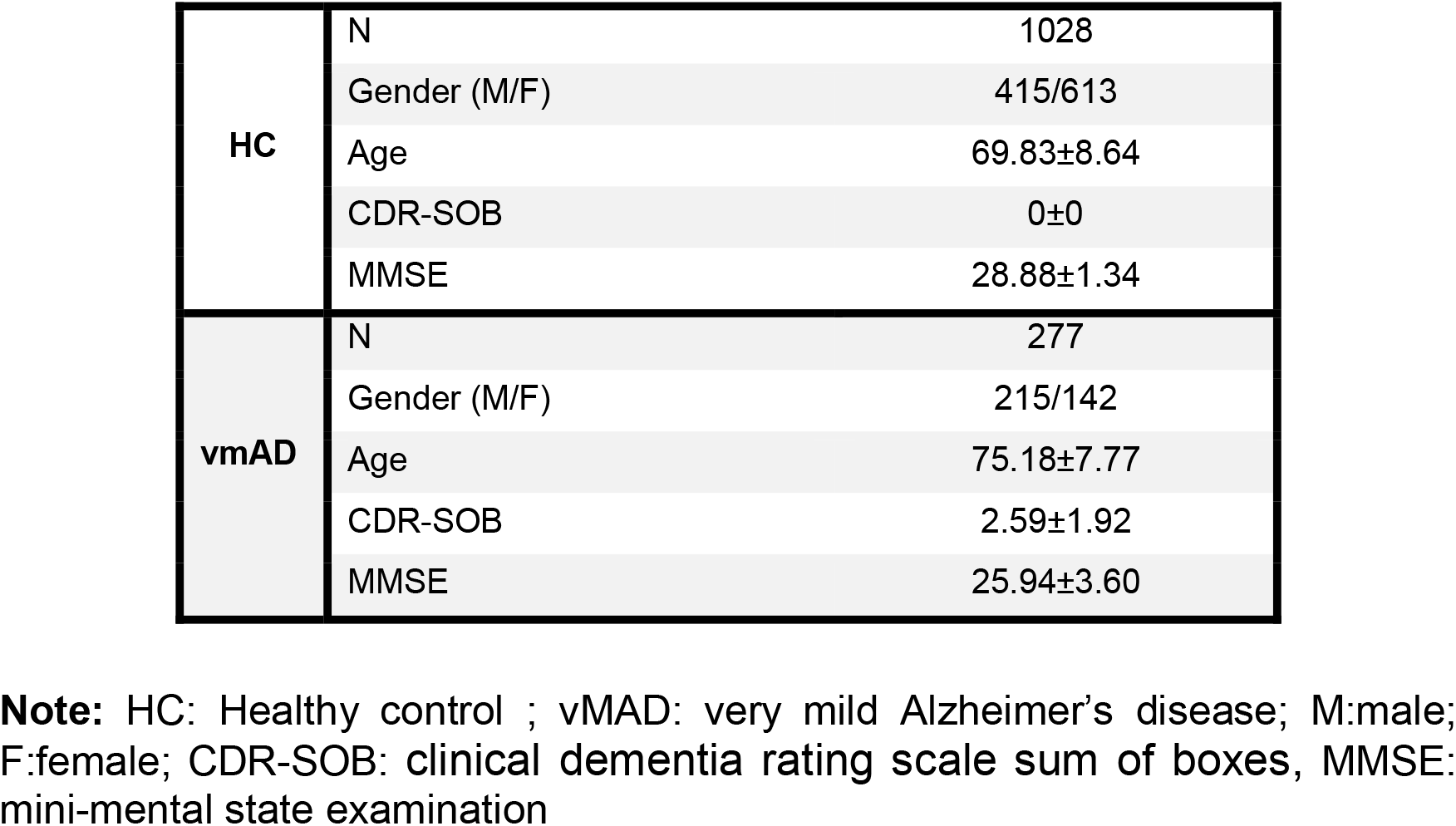
Demographic and clinical information.

|  |  |  |
| --- | --- | --- |
| **HC** | N | 1028 |
|  | Gender (M/F) | 415/613 |
|  | Age | 69.83±8.64 |
|  | CDR-SOB | 0±0 |
|  | MMSE | 28.88±1.34 |
| **vmAD** | N | 277 |
|  | Gender (M/F) | 215/142 |
|  | Age | 75.18±7.77 |
|  | CDR-SOB | 2.59±1.92 |
|  | MMSE | 25.94±3.60 |

**Note:** HC: Healthy control ; vmAD: very mild Alzheimer's disease; M: male; F: female; CDR-SOB: clinical dementia rating scale sum of boxes, MMSE: mini-mental state examination

### Data Acquisition

MRI data were collected from two scanners of TIM Trio 3T with a 20-channel head coil and foam pad stabilizers placed next to the ears to decrease motion (Siemens Medical Solutions USA, Inc). High resolution T2*-weighted functional images were acquired using echoplanar imaging or EP sequence with TE =27 ms, TR = 2.2 s, flip angle = 90°, slice thickness = 4mm, slice gap (center-to-center) = 4 mm, matrix size = 64, and field of view (FOV)= 256×256×128 mm^3^. The duration of the scanning was 6 minutes.

### Data processing

We processed the fMRI data using statistical parametric mapping (SPM12, https://www.fil.ion.ucl.ac.uk/spm/) in the MATLAB2019 environment. The first five dummy scans were discarded before preprocessing, and a slice-timing correction was performed on the fMRI data. To account for the subject’s head motion, we used rigid body motion correction. Next, the imaging data underwent spatial normalization to echo-planar imaging (EPI) template in the standard Montreal Neurological Institute (MNI) space and was resampled to 3×3×3 mm^3^. Finally, we used a Gaussian kernel to smooth the fMRI images using a full width at half maximum (FWHM) of 6 mm.

We used the Neuromark automated independent component analysis (ICA) pipeline, which uses previously derived component (region) maps as priors for spatially constrained ICA (Du et al., 2019), to extract reliable CCN independent components (ICs) or components. In Neuromark, replicable components were identified by matching group-level spatial maps from two large-sample healthy control (HC) datasets. Components were identified as meaningful regions if they exhibited peak activations in the gray matter within CCN. Seventeen data-driven CCN components were identified. These components are shown in Table 2.

**Table 2.** Component Labels.

| **Component name** | **Peak coordinate (mm)** |  |  |
| --- | --- | --- | --- |
| (IC 68) Inferior parietal lobule [IPL] | 45.5 | -61.5 | 43.5 |
| (IC 33) Insula | -30.5 | 22.5 | -3.5 |
| (IC 43) Superior medial frontal gyrus [SMFG] | -0.5 | 50.5 | 29.5 |
| (IC 70) Inferior frontal gyrus [IFG] | -48.5 | 34.5 | -0.5 |
| (IC 61) Right inferior frontal gyrus [R IFG] | 53.5 | 22.5 | 13.5 |
| (IC 55) Middle frontal gyrus [MiFG] | -41.5 | 19.5 | 26.5 |
| (IC 63) Inferior parietal lobule [IPL] | -53.5 | -49.5 | 43.5 |
| (IC 79) Left inferior parietal lobule [L IPL] | 44.5 | -34.5 | 46.5 |
| (IC 84) Supplementary motor area [SMA] | -6.5 | 13.5 | 64.5 |
| (IC 96) Superior frontal gyrus [SFG] | -24.5 | 26.5 | 49.5 |
| (IC 88) Middle frontal gyrus [MiFG] | 30.5 | 41.5 | 28.5 |
| (IC 48) Hippocampus [HiPP] | 23.5 | -9.5 | -16.5 |
| (IC 81) Left inferior parietal lobule [L IPL] | 45.5 | -61.5 | 43.5 |
| (IC 37) Middle cingulate cortex [MCC] | -15.5 | 20.5 | 37.5 |
| (IC 67) Inferior frontal gyrus [IFG] | 39.5 | 44.5 | -0.5 |
| (IC 38) Middle frontal gyrus [MiFG] | -26.5 | 47.5 | 5.5 |
| (IC 83) Hippocampus [HiPP] | -24.5 | -36.5 | 1.5 |

### Dynamic functional connectivity (dFC)

For each subject, i = 1 … N, the dFC of the 17 CCN components was estimated via a sliding window approach, as shown in Fig. 1. A tapered window obtained by convolving a rectangle (window size = 20 TRs = 44 s) with a Gaussian (σ = 3 s) was used to localize the dataset at each time point. A correlation matrix, based on Pearson correlation, was calculated to measure the dFC between CCN components (Step 1 in Fig. 1). The dFC estimates of each window for each subject were concatenated to form a (C × C × T) array (where C=17 denotes the number of components and T=139 windows), which represented the changes in brain connectivity between CCN components as a function of time (Calhoun et al., 2014).

**Fig 1.**
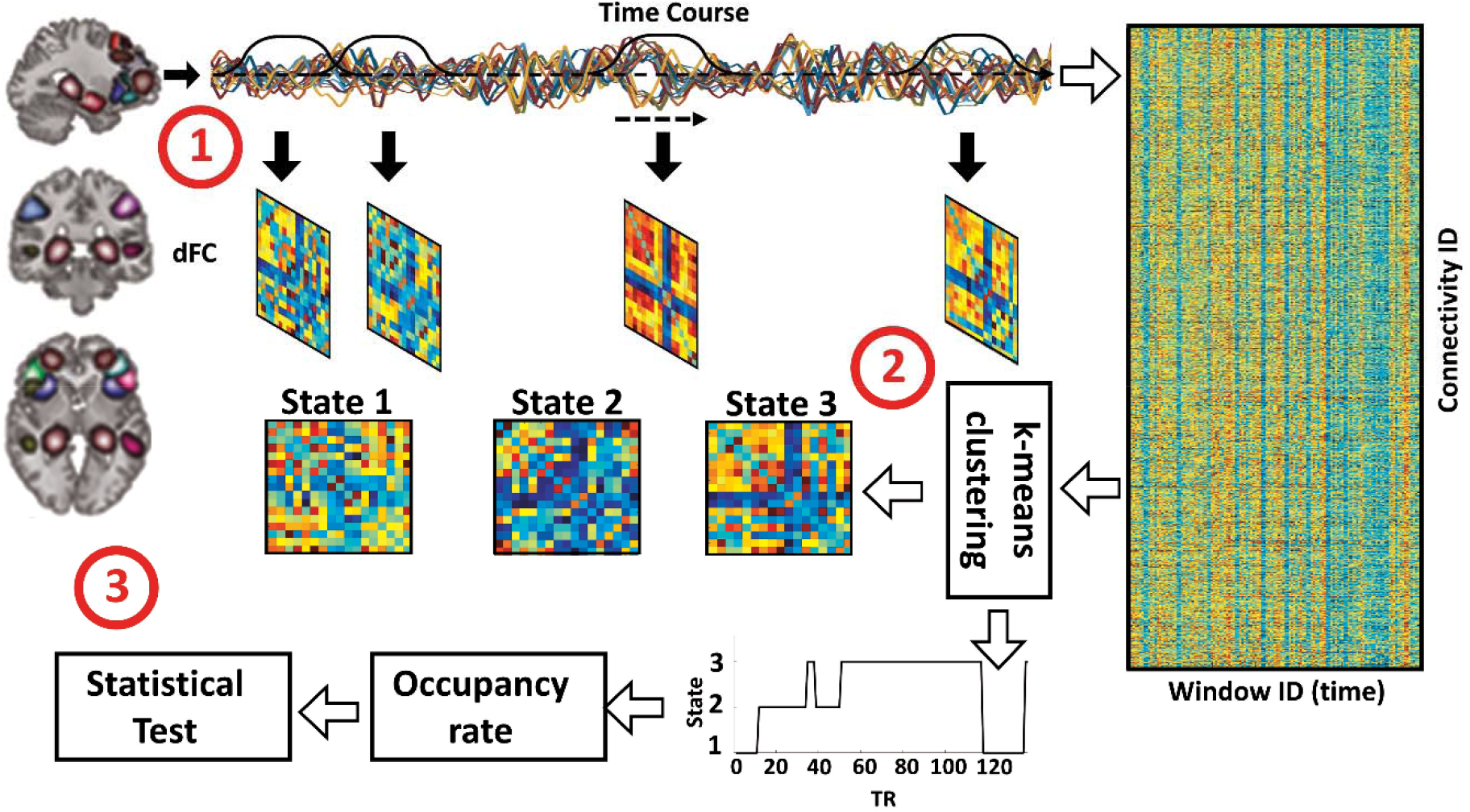
Analytic pipeline. Step1: The time-course signal of 17 components in the cognitive control network (CCN) has been identified using group-ICA. After identifying 17 regions in CCN, a taper sliding window was used to segment the time-course signals and then calculated the functional connectivity (FC). Step2: After vectorizing the FC matrixes, we have concatenated them, and then a k-means clustering, k=3, was used to group FCs to three distinct states. Elbow criteria were used to find the optimal k. In addition, the Euclidean distance metric is used in this clustering. Step3: Then, based on the state vector of each subject, the occupancy rate or OCR features, in total 3 features, were calculated from the state vector of each subject. Then, we explored the link between the state-specific connectivity feature and the clinical rate. In addition, we investigated the link between OCR with the clinical rate.

### Clustering and Latent Feature Estimation

A k-means algorithm was applied to the dFC windows to partition the data into a set of separated clusters. The optimal number of centroid states was estimated to be 3 using the elbow criterion based on the ratio of within to between cluster distance. This optimal number was found by searching between 2 to 9. We used the Euclidean distance metric in the clustering algorithm with 1000 iterations (Allen et al., 2014; Calhoun et al., 2014). The clustering output was 3 states for all subjects and state vector for each subject (Step 2 in Fig. 1). Next, to model the temporal changes of dFC for each subject using the state vector, we computed the OCR of dFCs in each state. OCR represents the proportion of time each subject occupies a given state. (Step3 in Fig. 1). From 3 states, we calculated 3 OCR features for each subject.

### Statistical Analysis

To find a link between dFC features, including OCR, and the cell values of the dFC in each state with mini-mental state examination (MMSE) score, we used partial correlation accounting for age and gender. We performed statistical analysis on the link between 136 state-specific CCN features obtained from 17 CCN components of each subject and clinical score, and between 3 OCR features obtained from state vectors of each subject and the clinical score, separately. All p values (3 p-values for the OCR and 136 p-values for state-specific CCN features with clinical score) have been adjusted by the Benjamini-Hochberg method for false discovery rate or FDR (Yoav Benjamini □; Yosef Hochberg, 1995).

## 3 Results

### Clinical and demographic results

The mean CDR-SOB of HC and vmAD were 0±0, 2.59±1.92, respectively. The mean MMSE of HC and vmAD were 28.88±1.34, 25.94±3.60, respectively. Using a two-sample t-test, we found significant differences in MMSE between HC and vmAD (p<1e^-5^). The mean age of HC and vmAD were 69.83±8.64 and 75.18±7.77, respectively. Using the two-sample t-test, we found significant differences between HC and vmAD age (t(1384)= −10.62, uncorrected p= 3.91e^-24^). It worth noticing that due to the significant age differences among the groups, we controlled the age and gender in all correlation analysis we did in this study.

### Overview of dFC states

Fig. 2 shows the 3 reoccurring dFC states, including state 1, state 2, and state 3 identified by k-means clustering. In all states, FCs within inferior parietal lobule, inferior frontal gyrus, the middle frontal gyrus, and the hippocampus were positive. In all states, among all subregions of CCN, the hippocampus had the lowest connectivity with the rest of CCN. Also, among all states, state 1 showed the strongest negative connectivity between the hippocampus and the rest of CCN. In addition, state 2 showed the least within the middle frontal gyrus comparing with that of other states. This state relatively showed more connectivity between hippocampus and the rest of CCN compared with that of other states. Also, state 3 showed the strongest positive connectivity within the inferior frontal gyrus. Finally, we measured the OCR of each subject in state1, state2, and state 3. OCR represents the amount of time each subject spends in each state. We found subjects spend 33.68 %, 28.02 %, and 38.30 % in state 1, state 2, and state 3, respectively. In addition, using a paired t-test, we found a significant difference between the OCR of state 1 and state 2 (t(1384)= 4.62, FDR corrected<1e^-4^) and between state 1 and state 3 (t(1384)= 3.73, FDR corrected p<1e^-4^), and between state 2 and state 3 (t(1384)= 8.62, FDR corrected p<1e^-4^).

**Fig 2.**
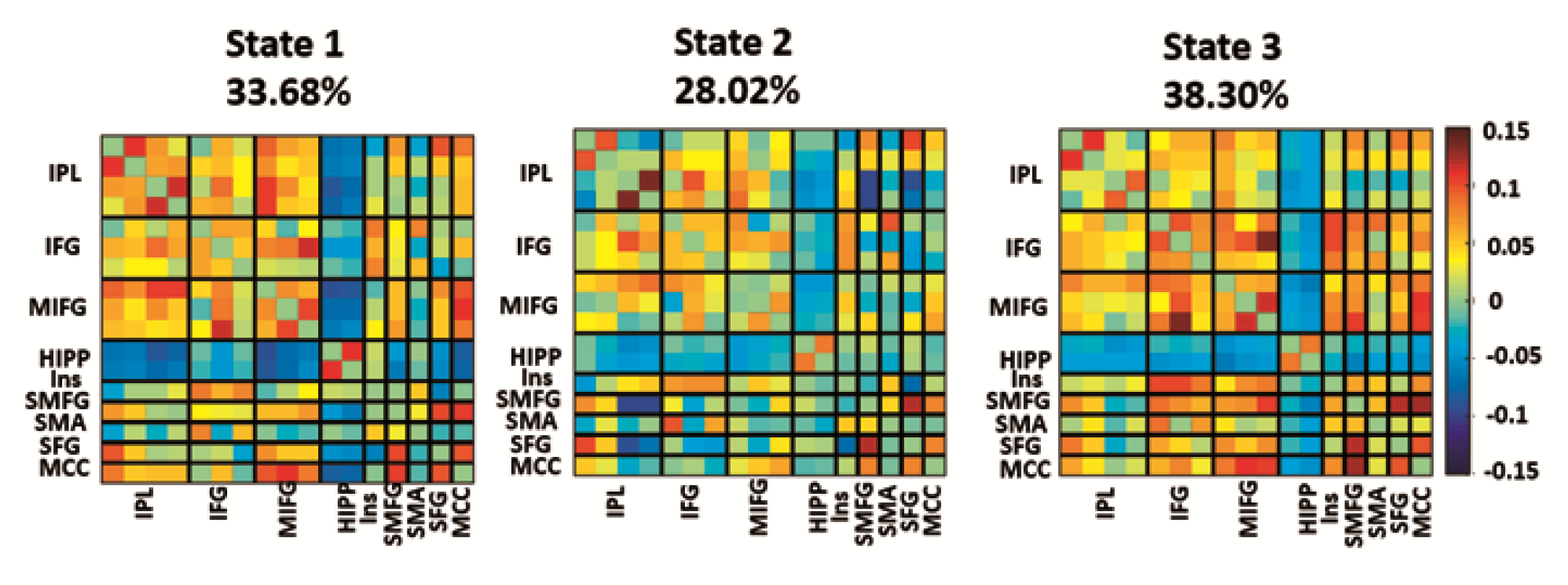
Dynamic functional connectivity (dFC) states. The three identified dFC states using the k-means clustering method. Each state is a 17×17 matrix in which the positive connectivity is shown with hot, and the negative connectivity is shown in cold color. We put all 17 components in 9 regions including inferior parietal lobule (IPL), inferior frontal gyrus (IFG), middle frontal gyrus (MIFG) hippocampus (HIPP), insula (INS), Superior medial frontal gyrus (SMFG), supplementary motor area (SMA), superior frontal gyrus(SFG), and middle cingulate cortex (MCC). In all states, the hippocampus showed a connection with the rest of the CCN. However, the connectivity between the hippocampus and the rest of the CCN in higher in state 2. Also, state 3 showed the strongest positive connectivity within the inferior frontal gyrus. We found subjects spend 33.68 %, 28.02 %, and 38.30 % in state 1, state 2, and state 3, respectively.

### Correlation between dFC cell features and MMSE

In each state, we averaged the connectivity features of all dFC for each subject. In more detail, each subject has multiple dFC in each state. Then, in each state, we used the average of dFC features (i.e., the average of 136 connectivity features) of each subject as her/his state-specific FC. Then, we calculated the partial correlation between averaged cell features of each subject and MMSE by controlling the age and gender to explore how these features changed by progressing from HC stages to AD stages. These results are shown in Fig. 3. The promising correlations (FDR uncorrected p<0.05) are shown in red (positive correlation) and blue (negative correlation). Also, a significant correlation that passes the FDR correction is marked by asterisks.

**Fig 3:**
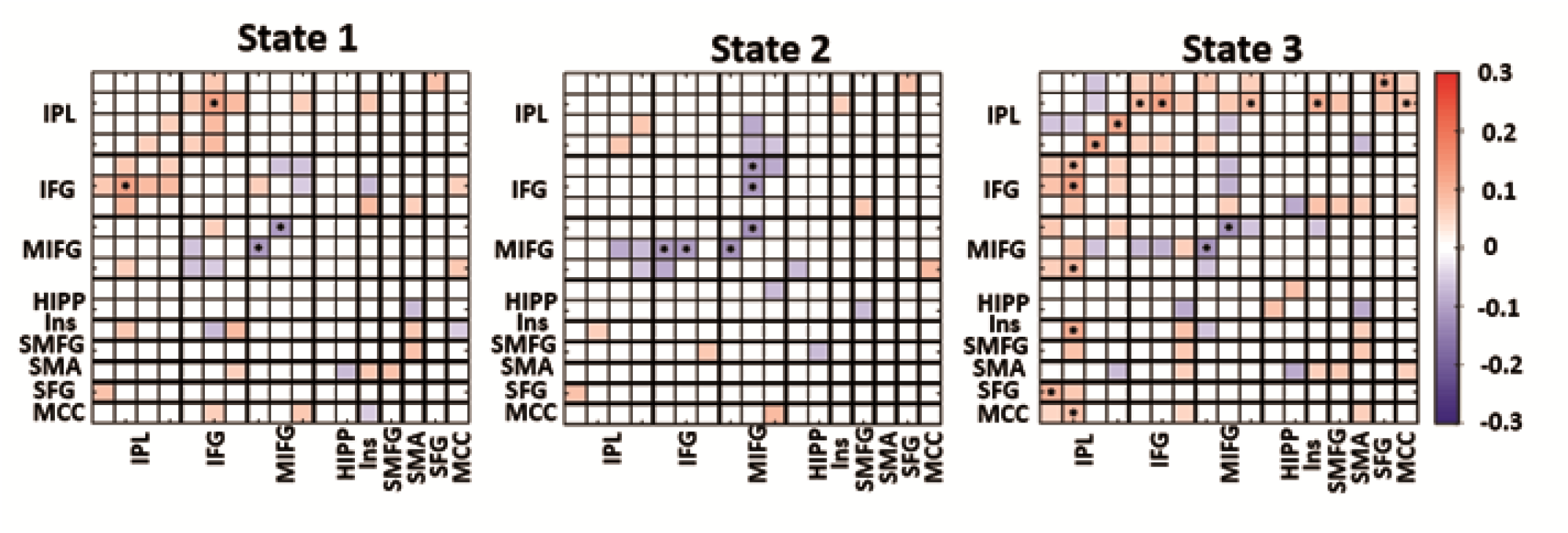
Correlation between FC of each state and MMSE. In each state, we averaged the cell features of all dFC of each subject, and then we calculated the partial correlation between averaged cell features of each subject and MMSE by controlling the age and gender to explore how these features changed by progressing from normal stage to AD stages. The correlation with significant p value (p ≤0.05) is highlighted in red and blue. A significant group difference that passes the multiple comparisons is marked by asterisks (false discovery rate [FDR] corrected, q = 0.05). Inferior parietal lobule (IPL), inferior frontal gyrus (IFG), middle frontal gyrus (MIFG) hippocampus (HIPP), insula (INS), Superior medial frontal gyrus (SMFG), supplementary motor area (SMA), superior frontal gyrus(SFG), and middle cingulate cortex

**Fig 4:**
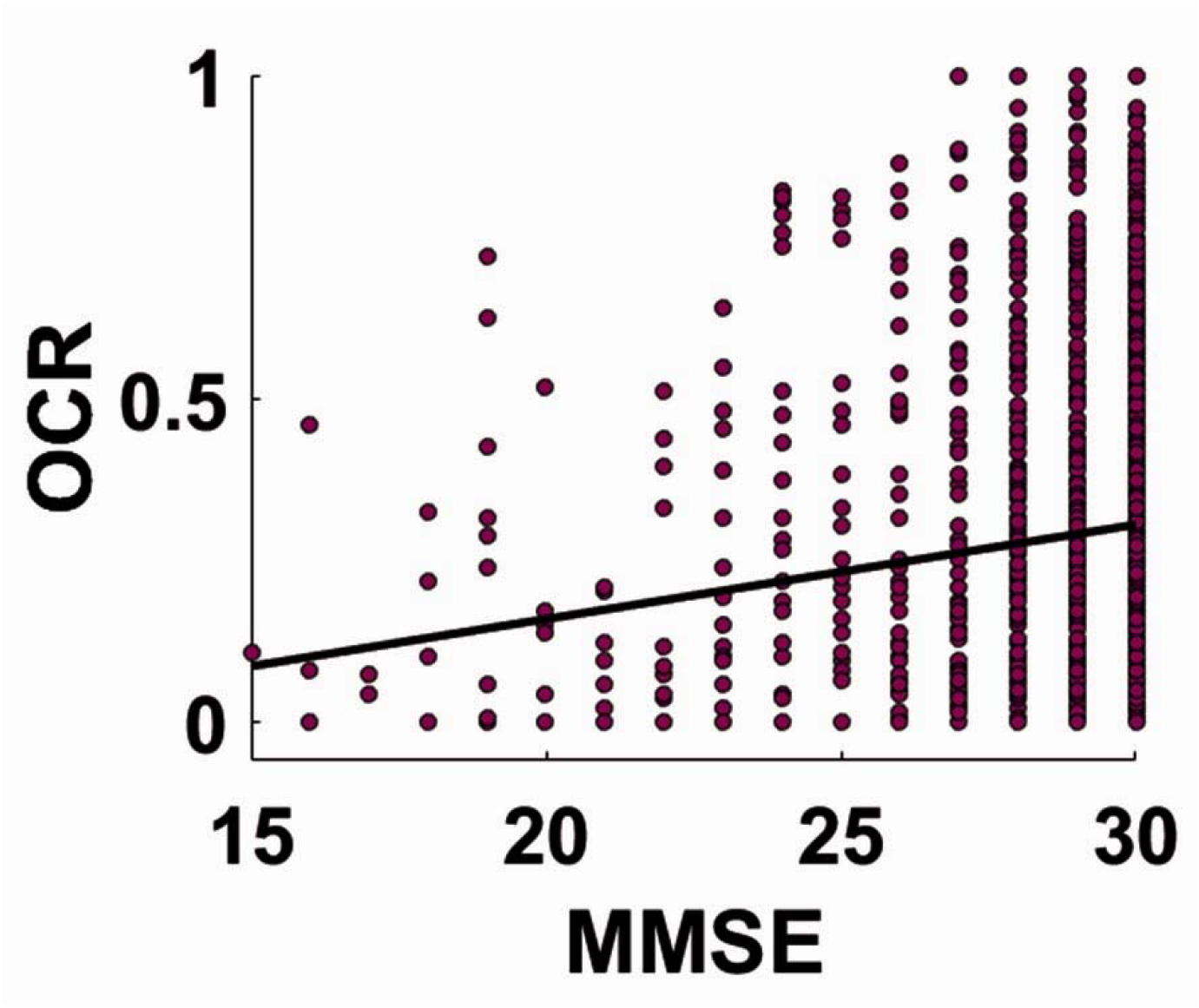
Correlation between OCR and MMSE. The partial correlation between the occupancy rate of state 2 with MMSE. r=0.073, FDR corrected p= 0.01, n=1385.

In state1, we observed a positive correlation between MMSE and the connectivity of inferior parietal lobule connection with inferior frontal gyrus. Also, we found a negative correlation between MMSE and the connectivity within the middle frontal gyrus. In state 2, we found that within middle frontal gyrus connectivity and its connectivity with inferior frontal gyrus showed a negative link with MMSE. In state 3, we found that the connection between inferior frontal lobule and the rest of CCN, including inferior frontal gyrus, middle frontal gyrus, insula, superior frontal gyrus, and middle cingula cortex showed a positive correlation with MMSE. Similar to state 1 and state 2, we observed a negative link between within middle frontal gyrus connectivity and MMSE.

### Correlation between dFC temporal pattern and MMSE

To find a link between MMSE and temporal features of dFC, including OCR, we used partial correlation by controlling age and gender. By calculating the correlation between MMSE and OCR, we found the OCR of state2, which representing spending time in state 2, shows a positive correlation with MMSE (Fig. 5: r=0.073, FDR corrected p= 0.01, n=1385). This means subjects with more severe symptoms spent less time in state 2, which showed the least middle frontal gyrus connectivity.

## 4 Discussion

Recent studies proved that the brain functional connectivity obtained from rs-fMRI is exceedingly dynamic and can disclose the underlying mechanism differences of brain connectivity in many diseases groups (Damaraju et al., 2014; Dong et al., 2019; Fiorenzato et al., 2019; Fu et al., 2019; Garrity et al., 2007; Miller et al., 2016; Sun et al., 2019; Zhi et al., 2018). In addition, it seems that the cognitive control network (CNN), which plays a significant role in several cognitive functions, degrades as AD progress (Dhanjal and Wise, 2014). Therefore, in the current study, we posit that studying the spatiotemporal pattern of CCN functional connectivity would add new information about the progression of AD. In more detail, we investigated the dFC of several data-driven components of CCN, including inferior parietal lobule, inferior frontal gyrus, middle frontal gyrus, hippocampus, insula, middle cingula cortex, and superior frontal gyrus. We found that functional connectivity in CCN is indeed highly dynamic, representing flexibility in functional coordination in this mode. Then, in each state, we used a partial correlation between FC features and MMSE to explore whether the FC features show a link with cognitive scores or not.

We found that the connectivity of the inferior parietal lobule with other regions of CCN has a positive correlation with MMSE in state 3, which means by progression from the normal state to AD state, the FC of the inferior parietal lobule with other regions decreases. Some evidence from neuroimaging studies implies the role of the inferior parietal lobule in a broad range of behavior and function, including cognitive functionality (Bzdok et al., 2016; Caspers et al., 2013; Wang et al., 2017). For example, a previous study proved the vital role of an inferior parietal region in maintaining attentive control (Shapiro and Hillstrom, 2002). A reduction of gray matter volume in the inferior parietal lobule by the progression of AD is reported (Greene and Killiany, 2010; Oishi et al., 2018). A recent research paper reported an improvement of working memory in AD patients by applying transcranial direct current stimulation to the inferior parietal region (Roncero et al., 2017). All these studies emphasized the role of this region in AD progression. However, inferior parietal lobule functional connectivity is understudied.

In our study, using a relatively large dataset, we found that inferior parietal lobule functional connectivity is affected as AD progresses, more so than other regions of CNN studied here, as well as in the early stage of AD by progression from the normal brain to AD. Therefore, this region could be a reasonable brain area to target for future AD studies. However, this connectivity does not show a significant correlation with MMSE in state 2. This aberrant spatiotemporal pattern in the connectivity of inferior parietal lobule and CCN subregions potentially underlined the importance of studying the dFC and analyzing the FC in a shorter period of time. A recent study showed a disrupted pattern between the inferior parietal lobule and default mode, salience, executive control, and sensorimotor networks (Wang et al., 2015). The current study provides new knowledge about the disrupted pattern between the inferior parietal lobule and the rest of CCN.

The inferior frontal gyrus also showed a disrupted pattern. Although we found that the connectivity between inferior frontal gyrus and inferior parietal lobule in state 1 and state 3 decreases by progression from healthy to AD, the connectivity between inferior frontal gyrus and middle frontal gyrus of state 2 increases in this progression. In a pilot study, the inferior frontal gyrus was stimulated by transcranial magnetic stimulation for improving attention in the early stage of AD. However, the result was not consistent and significant in all tests (Eliasova et al., 2014). Our result of having multiple distinct patterns in the inferior frontal gyrus connectivity further highlights the importance of additional studies evaluating the potential of AD intervention by modulating these connectivity patterns.

Interestingly, we found that the increase within middle frontal gyrus connectivity positively correlates with cognition decline in AD with mild symptoms. A previous study introduced a compensatory mechanism of neural resources in AD by showing more activation of the middle frontal gyrus in AD patients introduced as (Woodard et al., 1998). In the current study, a higher middle frontal gyrus connectivity in AD would further support the compensatory mechanism to reduce the impact of the decrease of connectivity in other regions of CCN in AD subjects (Gaubert et al., 2019). Also, like the inferior parietal lobule and inferior frontal gyrus, we observed a disrupted pattern in the connectivity between the middle frontal gyrus and CCN subregions. Interestingly, the hippocampus is showing the least effect among all regions in the CCN. This result is consistent with a previous study showing that the hippocampus connectivity shows fewer changes in early-onset AD (Park et al., 2017). Compared with other areas in CCN, these pieces of evidence might suggest that hippocampus connectivity with other subregions of CCN is one of the last brain areas affected by AD. However, a previous study found a dysconnectivity between the hippocampus and posterior cingulate cortex (PCC) as a part of DMN (Grieder et al., 2018). Therefore, our finding might be limited to the connectivity between hippocampus and CCN subregions, and the connectivity between the hippocampus and other brain networks beyond CCN, might be disrupted with AD progression.

Next, to model the temporal pattern of dFC, we estimated the OCR. Also, to explore how the temporal pattern of dFC correlates with cognition, we calculated the correlation between OCR and MMSE by controlling age and gender. In this analysis, we found dwell time of state 2 showed a significant and positive correlation MMSE. This means that subjects participants with mild impairment and mild symptoms tend to stay less in a state, which showed the least connectivity within the middle frontal gyrus among all states, than a normal brain. This provides further evidence of the effect of disease on the dysregulating temporal properties of CCN FC.

In addition, state 2 showed relatively higher connectivity between the hippocampus and the rest of the brain. Although we did not find any significant correlation between hippocampus connectivity strength and clinical rate, the temporal dysregulation effect of the AD progression might reveal some new information about the effect of this disease on the hippocampus, even in the early stage of the disease. This finding potentially highlighted the importance of the study of functional connectivity in a shorter period. Previous studies highlighted the effect of AD on the temporal pattern on brain FC. One study with 29 AD patients and 31 HC subjects found that AD patients spend more time than the HC subject in a state with sparse connectivity pattern in which the motor network is isolated from the rest of the brain. In addition, the same study found an inability to switch out from a state with low inter-network connectivity into more highly connected network configurations in AD patients (Schumacher et al., 2019). In another study, an altered dFC temporal pattern has been shown in Parkinson’s disease with dementia, consistent with the result of the current study (Fiorenzato et al., 2019). In the current study, we provided new evidence about the effect of AD progression on altering the temporal pattern of CCN.

### Limitations and Future directions

There are some limitations to this work. MMSE is commonly used to measure the cognitive state of the brain in the different stages of AD. However, a few studies suggested that MMSE is a noisy measure in the diagnosis of AD (Wang et al., 2015). The choice of window size is an implicit assumption about the dynamic behavior in that a short window captures more rapid fluctuations, whereas a longer window does more smoothing than a shorter one. Previous dFC studies showed that the window size between 30-60 s is a reasonable choice for capturing dFC fluctuation and any widow size above the safety limit, i.e., the largest wavelength present in the preprocessed fMRI time courses, would not change the result significantly (Preti et al., 2017). In addition, k-means clustering needs predefined knowledge for setting the clustering parameters, including the distance metrics. By applying different distance metrics, we did not observe a significant difference in the results, however applying other clustering methods such as robust continuous clustering would eliminate this shortcoming and might improve the clustering results (Shah and Koltun, 2017).

As mentioned earlier, a previous study showed a dysconnectivity between the hippocampus as a part of CCN and DMN in AD (Grieder et al., 2018). Therefore, a prospective study on the effect of this dysconnectivity between CCN and DMN, and its links with AD progression is needed. Although in the current study we focused on the CCN based on the prior knowledge of its role in cognitive impairment, future studies and methods that can mechanistically remove the irrelevant networks in study dynamic functional connectivity are needed (Cohen et al., 2015; Qiao et al., 2019; Schlesinger et al., 2017).

### Conclusion

To summarize, we explored the alteration of temporal features of CCN FC by comparing participants of the normal state, mild impairment, and mild dementia. We found that AD progression affects differently on CCN subregions. It decreases inferior parietal lobule connectivity with other regions and increases within-middle frontal gyrus connectivity. We found both decrease and increase in inferior frontal gyrus and middle frontal gyrus connectivity with the rest of CCN subregions. Also, we found by progression from normal to mild dementia stage decreases the dwell time in a state with less within middle frontal gyrus and higher connectivity between the hippocampus and rest of the CCN. This result further supports the role of the middle frontal gyrus in the progression of AD. Interestingly, while local (cell-wise) hippocampal patterns were not impacted by AD progression, more global (state-based) patterns linking CCN to hippocampus showed lower occupancy as AD progressed. Our results suggest that AD progress changes not only the connectivity strength but also affects the temporal properties in the brain network connectivity. In other words, our finding posits the temporal and spatial pattern of CCN as a biomarker in AD that differentiates patients based on symptom severity.

## Author Contribution

Mohammad S. E. Sendi developed the study, conducted data analysis, interpreted the results, and wrote the original manuscript draft. Elaheh Zendehrouh conducted data analysis. Zening Fu conducted data analysis. Jingyu Liu edited the original draft and provided critical review to the initial draft. Yuhui Du developed the Neuromark and provided critical review to the initial draft. Elizabeth Mormino provided critical review to the initial draft. David H. Salat edited the original draft and provided critical review to the initial draft. Vince D. Calhoun developed the study, interpreted the results, edited the original draft, and provided critical review to the initial draft. Robyn L. Miller developed the study, interpreted the results, edited the original draft, and provided critical review to the initial draft. All authors approved the final manuscript.

## Acknowledgement

We thank those who helped collect this valuable data.

## Abbreviations

AD: Alzheimer’s disease
vmAD: very mild Alzheimer’s disease
dFC: Dynamic functional connectivity
sFC: Static functional connectivity
MCI: Mild cognitive impairment
CCN: Cognitive control network
OCR: Occupancy rate
CDR-SOB: Clinical dementia rating scale sum of boxes
HC: Healthy control
EPI: Echo-planar imaging
MNI: Montreal Neurological Institute
FWHM: Full width at half maximum
MMSE: Mini-mental state examination
ICA: Independent component analysis
IC: Independent component

